# Grip-preference distribution among Asian-Oceanian floorball players is opposite that of European and North American players – How environmental constraints may influence human motor behaviour

**DOI:** 10.1101/2022.11.10.515948

**Authors:** Karen Emilia Ekman, Arve Vorland Pedersen

**Affiliations:** Department of Neuromedicine and Movement Science, Faculty of Medicine and Health Sciences, Norwegian University of Science and Technology, Trondheim, Norway

**Keywords:** motor learning, bimanual coordination, asymmetry

## Abstract

Whilst most humans are right-handed, handedness alone cannot explain the large variability observed in bimanual motor behaviour. Sport-specific motor behaviour provides a natural laboratory for laterality and motor-control research. It is known among floorball players, that Europeans more often play using a left-sided grip, whereas most Asian players are right-gripped, with no logical explanation. However, the exact grip-side distribution is unknown. The present study investigated the influence of environmental constraints on lateral motor behaviour by assessing geographic variabilities in floorball-specific grip preferences between European and Asian national team floorball players.

A small-scale Big Data approach was utilised to collect data on lateral preferences for both field players and goalkeepers from the International Floorball Federation website. Data included 2,935 players representing 40 national teams from three different confederations (Europe, Asia-Oceania, and North America). More than two-thirds of European and North American players preferred a left-sided grip, whereas the same number of Asian-Oceanian players preferred a right-sided grip.

To the best of our knowledge, these are the first findings of such large geographic variations in any lateralised motor behaviour. No biological factors are likely responsible for the difference in lateral-preference distribution. Environmental and task-specific constraints are discussed as possible explanations.

**Summary Box**
Sport-specific laterality is generally believed to be correlated with general handedness. In floorball, the bimanual grip-preference distribution is the exact opposite for European versus Asian-Oceanian national team field players, whilst goalkeepers’ throwing-hand preference is similar across all confederations and, thus, aligned with general handedness. Lateral preferences in a sport may develop somewhat independently of athletes’ general handedness, possibly guided by external environmental factors.

## Introduction

It is generally known that 90% of the general population prefer to use their right hand for most manual activities. Less is known about the large intra-individual variability across different tasks (1, 2) or the large inter-individual variability with respect to both direction and degree of manual preferences (3). Knowledge about the sometimes systematically skewed distributions of lateral preferences within certain populations due to external variables that can be beyond the control of human intention is particularly lacking. For example, among top performers within many sports, the prevalence of left-handers is elevated, not because they are selected based on their handedness but because being left-handed is an advantage (4, 5). This advantage can be explained by frequency dependence; that is, those who are in the minority have an advantage in combat with those representing the majority. This is referred to as the ‘fighting hypothesis’ (4, 6, 7).

The predominant lateral preference in most unimanual sports is, similar to global general handedness, rightward (4, 5). However, despite the two being correlated, the variability is large across sports and across various actions within each sport, and thus, sport-specific lateral preferences may not reflect an individual’s general laterality (4, 8). The correlation is particularly weak when assessing lateral preferences for bimanual actions (4). For example, a right-sided grip preference has been shown to constitute the majority laterality in sports, such as batting in baseball and cricket and swinging a golf club (9-12). However, the opposite pattern is evident in ice hockey (13).

Knowing what makes a variable (in this case, lateral preference) vary and the extent to which it is important, of course, for being able to make hypotheses about that variable or for basing decisions on it (for example, when designing interventions or research studies). Thus, which of the independent variables acts on the dependent variable and in what way must be determined.

In motor-behaviour research, such independent variables are often referred to as constraints on human movement. This was specifically described by Newell (1986), who distinguished between *individual constraints* (within the individual, e.g. height, weight and body composition), *task constraints* (within the task being performed, e.g. the goal, rules and equipment) and *environmental constraints* (general or task-specific properties of the environment, e.g. gravity, temperature, light and obstacles) (14). The third category, it has been argued, would also include social and cultural constraints (15). Situation-specific constraints create competing boundaries, according to which a functional coordination pattern emerges in individual performers (14).

Due to the stable ratio of right- and left-handedness throughout history, handedness has generally been believed to be heritable and, thus, caused by *individual constraints*. However, heritability has been reported to explain about only 25% of the factors that determine an individual’s laterality, leaving 75% to non-genetic factors (16). Some ancestral (geographic) variations in left-handedness have been reported, with Europe having a higher prevalence (11%) compared to Africa (8%) and East Asia (6%) (5).

Situation-specific *task constraints* such as the aim of the task, sport rules or equipment have also been shown to shape an individual’s laterality. In sport-specific terms, the same baseball bat can be held with either a left-sided or right-sided grip, whereas the angle of the blade on an ice-hockey stick affords one laterality over the other.

Structures or features of the *environment* in which the sport or task takes place may favour one laterality over the other and, thereby, guide preferences or performance. For example, inside an ice-hockey rink, a player’s relation to the rink has been shown to influence performance. More specifically, left-sided players benefit from playing on the right side of the field and positioning their stick on the inside; the opposite is the case for right-sided players. Furthermore, a player’s relation to other dynamic structures (notably, other players) in the environment has also been shown to influence playing behaviour. For example, it has been suggested that, regardless of the goal scorer’s grip preference, more goals are assisted by left-sided players than by right-sided players, relative to their distribution (13). In regard to general handedness, the social environment has been shown to have the strongest constraining effect (due to stigmatisation of left-handers; see, for example, Kushner (17)).

Studying real-life variability often requires methods that differ distinctly from those applied to experimental designs or controlled trials, in that these are usually designed to reduce variability in order to make variables controllable. Sports is one context in which an abundance of variability can be found, and vast amounts of recorded data are easily accessible. Thus, as Furley (18) argued, sport provides a plethora of naturally occurring data that can be exploited to increase our understanding of human motor behaviour.

Floorball is a relatively new sport, but due to its immense growth in popularity, it is already regarded as a global, elite sport requiring advanced skills and great commitment (19, 20) and is recognised as such by the International Olympic Committee (21). Floorball is played indoors and resembles indoor field hockey or ice hockey. The field players play with plastic sticks with the aim of hitting a lightweight plastic ball into the opponent’s goal to score (22).

A floorball field player’s sport-specific laterality is defined according to the individual preference for holding the stick on either side of the body, referred to as left-sided or right-sided grip (see Figure 1). Left-siders will hold the upper part of the stick with the right hand, whereas right-siders will do so with their left hand. It is generally acknowledged as preferable, and thus advised by coaches and others, to place one’s dominant hand on the upper part of the stick (20).

**Figure 1.**
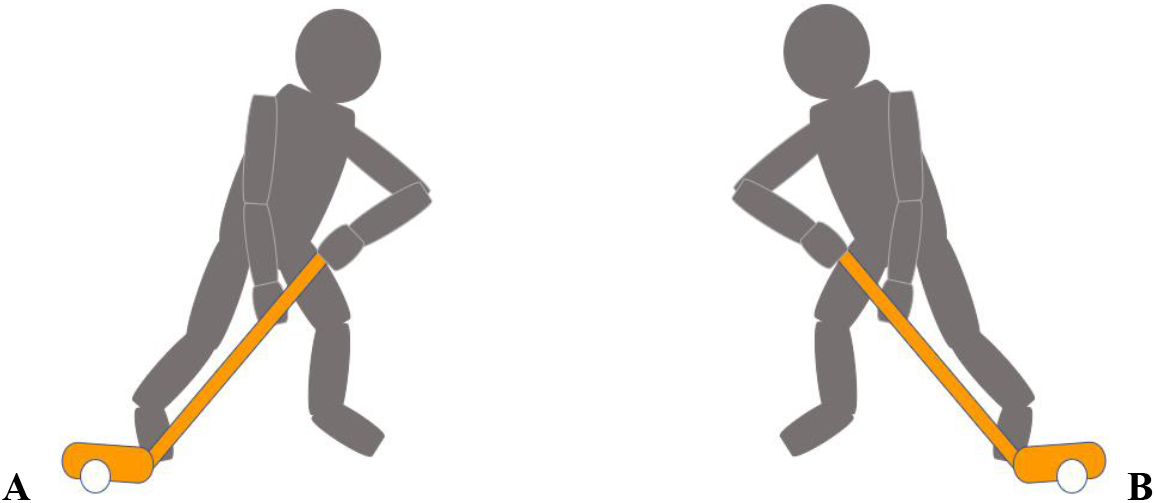
Right-sided (A) and left-sided (B) grip preferences. In a right-sided grip preference the player places the left hand at the upper part of the stick and the right hand towards the middle of the shaft, whereas a left-sided player places the right hand at the upper part and the left hand towards the middle

A left-sided grip is, however, preferred by the majority of European floorball players, which may be explained by the fact that the vast majority of the general population is right-handed (22). By contrast, it has been generally noted within the sport that the majority of Asian national team floorball players seems to prefer a right-sided grip (20, 22).

Floorball goalkeepers do not play with a stick in their hand (as they would in field hockey or ice hockey), thus freeing both hands for interceptive actions when guarding the goal and for initiating play after gaining possession of the ball. The main lateralised movement pattern of goalkeepers is the unimanual throw of the ball for initiating play (23), thus indicating that their hand-preference distribution would be expected to mimic that of the general population with a 90/10 split, as would be found in the handedness literature for throwing (24).

Such a phenomenon of completely opposite lateral motor behaviours between two different geographic areas, as described above, would indicate that the difference was not due to individual constraints or to any known task constraints. The individual constraints affecting floorball players’ lateral preferences are the same across the different confederations (5), as are the (floorball-specific) task constraints. Thus, the inverse preference distribution between confederations must be due to variations within one or more environmental constraints. If such a geographic variability is prevalent, it could act as proof of the strength with which environmental constraints might modify individual preferences and influence the emergence of functional motor behaviour.

The aim of the present study was to investigate the influence that environmental constraints may have on lateral motor behaviour. This was done by assessing geographic variabilities within bimanual motor behaviour and, specifically, by comparing floorball-specific grip preferences across different geographic, sex, and age groups. The following main hypotheses were tested:

1. Grip-preference distribution is different between the European and Asian-Oceanian confederations in that European national team players more often play with a left grip, whereas Asian-Oceanian national team players more often prefer a right grip.
2. There is no difference in regard to hand preference among goalkeepers between confederations.
3. The grip-preference distribution will be similar across sex and age groups.

## Methods

### Data collection

The data were collected from the International Floorball Federation’s official website (https://floorball.sport) and comprised all players from the women’s, men’s, and respective juniors’ national teams. Players listed on the current roster at the time of data collection, as well as players listed on the roster for all rounds of the most recently played world championship (WC), were included in the study, and their age group, nationality, playing position and grip preference were recorded.

All data were publicly available and did not include any sensitive information. The data were treated anonymously in accordance with relevant guidelines and regulations.

A total of 3,166 players were initially included in the dataset. Players whose grip preference was not reported were excluded (n=27). Whenever a player appeared in both Under-19 (U19) and adult team rosters, the player’s data were removed from the adults’ tables to avoid counting it twice in the total (n=204). After the exclusion process, a total of 2,935 players were included in the data analysis.

Playing positions were categorised according to the classification on the IFF website (FW = forward, DF = defence, GK = goalkeeper). Grip side was similarly reported and recorded as either L = left side or R = right side. Similarly, goalkeepers’ hand preference was categorised as L = left hand or R = right hand.

Data were collected between 10 September 2020 and 24 November 2020.

### Statistical measures

Frequency distributions and respective percentages of lateral preferences within each floorball confederation are presented in Table 1. These were compared between confederations as well as between sex and age groups within each confederation. Chi-square statistical tests in SPSS (IBM Statistics, v.26) were conducted with the level of significance set at 5% (p<0.05). However, since the sample equals the total population (n=all), the use of inferential statistics may be redundant and should not be used solely when judging the accuracy of the findings (25).

**Table 1.** Field players’ and Goalkeepers’ task specific left- and right-sided preferences

|  |  | Women |  | WU19 |  | Men |  | MU19 |  | Total Female |  | Total Male |  | Total |  |
| --- | --- | --- | --- | --- | --- | --- | --- | --- | --- | --- | --- | --- | --- | --- | --- |
|  |  | L (%) | R (%) | L (%) | R (%) | L (%) | R (%) | L (%) | R (%) | L (%) | R (%) | L (%) | R (%) | L (%) | R (%) |
| **Europe** | FP | 261<br>(73.73) | 93<br>(26.27) | 373<br>(78.20) | 104<br>(21.80) | 393<br>(67.64) | 188<br>(32.36) | 368<br>(73.02) | 136<br>(26.98) | 634<br>(76.29) | 197<br>(23.71) | 761<br>(70.14) | 324<br>(29.86) | 1395<br>(72.81) | 521<br>(27.19) |
|  | GK | 3<br>(6.38) | 44<br>(93.62) | 11<br>(18.33) | 49<br>(81.67) | 16<br>(23.88) | 51<br>(76.12) | 16<br>(26.23) | 45<br>(73.77) | 14<br>(13.08) | 93<br>(86.92) | 32<br>(25.00) | 96<br>(75.00) | 46<br>(19.57) | 189<br>(80.43) |
| **Asia-Oceania** | FP | 58<br>(30.85) | 130<br>(69.15) | 6<br>(12.77) | 41<br>(87.23) | 93<br>(39.41) | 143<br>(60.59) | 19<br>(23.17) | 63<br>(76.83) | 64<br>(27.23) | 171<br>(72.77) | 112<br>(35.22) | 206<br>(64.78) | 176<br>(31.83) | 377<br>(68.17) |
|  | GK | 4<br>(15.38) | 22<br>(84.62) | 2<br>(40.00) | 3<br>(60.00) | 1<br>(3.57) | 27<br>(96.43) | 0<br>(0.00) | 8<br>(100.0) | 6<br>(19.35) | 25<br>(80.65) | 1<br>(2.78) | 35<br>(97.22) | 7<br>(10.45) | 60<br>(89.55) |
| **North America** | FP | 17<br>(62.96) | 10<br>(37.04) | 34<br>(73.91) | 12<br>(26.09) | 23<br>(56.10) | 18<br>(43.90) | 22<br>(62.86) | 13<br>(37.14) | 51<br>(69.86) | 22<br>(30.14) | 45<br>(59.21) | 31<br>(40.79) | 96<br>(64.43) | 53<br>(35.57) |
|  | GK | 0<br>(0.00) | 3<br>(100.00) | 0<br>(0.00) | 4<br>(100.00) | 1<br>(20.00) | 4<br>(80.00) | 1<br>(33.33) | 2<br>(66.67) | 0<br>(0.00) | 7<br>(100.00) | 2<br>(25.00) | 6<br>(75.00) | 2<br>(13.33) | 13<br>(86.67) |
| **Total** | FP | 336<br>(59.05) | 233<br>(40.95) | 413<br>(72.46) | 157<br>(27.54) | 509<br>(59.32) | 349<br>(40.68) | 409<br>(65.86) | 212<br>(34.14) | 749<br>(65.76) | 390<br>(34.24) | 918<br>(62.07) | 561<br>(37.93) | 1667<br>(63.67) | 951<br>(36.33) |
|  | GK | 7<br>(9.21) | 69<br>(90.79) | 13<br>(18.84) | 56<br>(81.16) | 18<br>(18.00) | 82<br>(82.00) | 17<br>(23.61) | 55<br>(76.39) | 20<br>(13.79) | 125<br>(86.21) | 35<br>(20.35) | 137<br>(79.65) | 55<br>(17.35) | 262<br>(82.65) |

**Note:** % represents proportion of respective sex total, specific to confederation and playing position. FP = Field player, GK = Goalkeeper. WU19 = Women’s U19 junior team, MU19 = Men’s U19 junior team. FPs’ grip-side distributions in the ‘Total’ (lowest) row are not representative of the findings since all data from the different confederations are combined.

### Sample data

The final sample included 2,935 players (43.7% female and 56.3% male) aged 15 to 55 (M 23, ± 5.6). Of these, 1,603 represented womens’ (W) and mens’ (M) adult national teams and 1,332 represented junior U19 (WU19 and MU19) national teams. A total of 2,618 players were field players (1,019 defenders and 1,599 forwards) and 317 were goalkeepers. Due to the many differences in task constraints across playing positions, goalkeepers were analysed as a separate group. Further sample statistics are shown in Table 1.

The players originated from 40 countries. Activities of the respective national teams are managed by three main floorball associations, the European, Asian-Oceanian and North American Floorball Confederation, and players were thus categorised accordingly in the data set (n = 2,151, 620 and 164, respectively).

## Results

### Field players

The distribution of grip-side preference among field players varied significantly across the different confederations (ꭓ^2^ = 309.954, *p* < .0001).

Of 1,915 European field players, almost three-quarters of the sample played with a left-sided grip. Similarly, two-thirds of the 149 players in the North American sample reported a left-sided grip preference. By contrast, of the 554 Asian-Oceanian field players, one-third of the sample preferred a left-sided grip (see Figure 2). The differences between confederations were consistent across sex and age groups, although with several variations (see Table 1).

**Figure 2.**
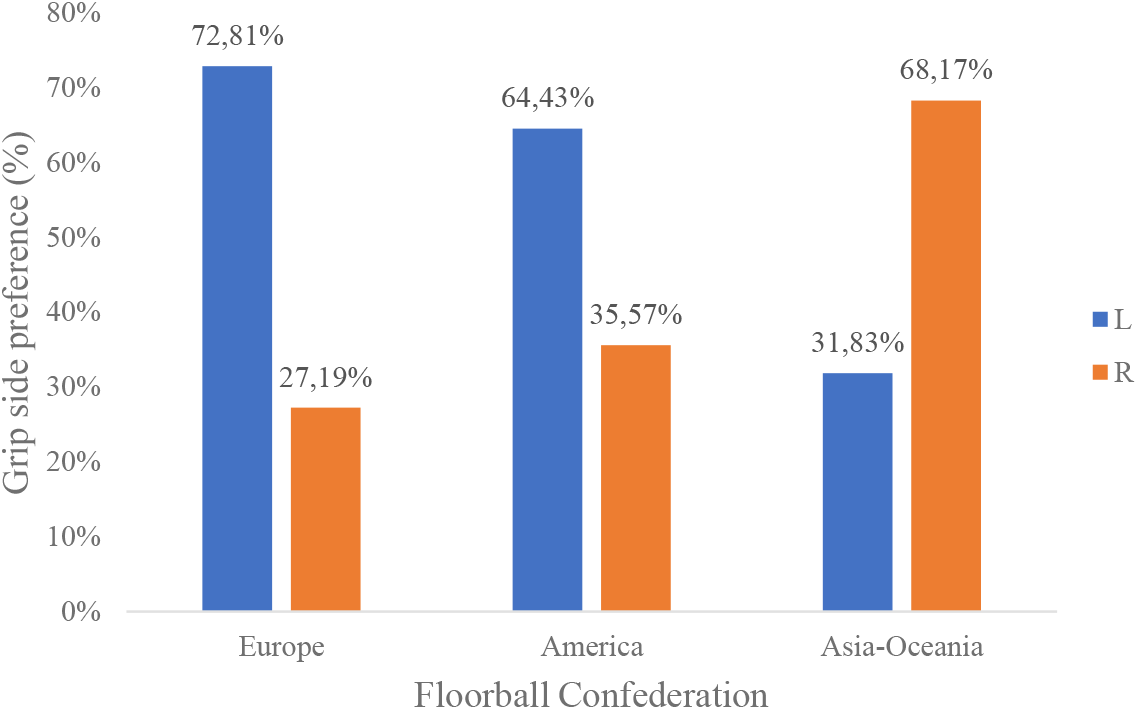
Proportion (%) of left- and right-sided grip preferences for field players in each floorball confederation

Variations in grip preference were found when comparing sexes. European men were more likely to prefer a (minority) right-sided grip than European women (ꭓ^2^ = 8.913, *p* = .003 <.05). Similarly, Asian men were somewhat more likely to prefer a (minority) left-sided grip than Asian women. The difference trended towards a significant level, although it did not meet the set standard of 5% (ꭓ^2^ = 3.673, *p* = .055 > .05). No significant difference between sexes in the North American confederation was found (ꭓ^2^ = 1.844, *p* = .175 > .05), but note the much smaller sample.

In the younger age groups (U19), there was a tendency for a higher prevalence of players to choose the confederation-specific majority laterality compared with their older counterparts on the senior teams. That is, European and North American U19 players were more often left-sided (75.5% and 69.1% respectively) compared to the players on the adult teams (69.9% and 58.8% respectively). The Asian-Oceanian U19 players exhibited a similar, although opposite, tendency for a higher prevalence of right-sidedness (80.6%) compared with the adult group (64.2%). The difference between age groups was significant in the Asian-Oceanian sample (ꭓ^2^= 12.220, *p* < .001 < .05) and the European sample (ꭓ^2^ = 7.633, *p* = .006 < .05) but not in the North American sample (ꭓ^2^ = 1.715, *p* = .190 > .05). Note, again, the smaller North American sample.

### Goalkeepers

Contrary to the findings for field players, no significant geographic variation was found in the reported (throwing) hand preference for floorball goalkeepers (ꭓ^2^ = 3.206, *p* = .203>.05). A right-hand preference was reported by the majority of all European, Asian-Oceanian and North American goalkeepers (see Figure 3).

**Figure 3.**
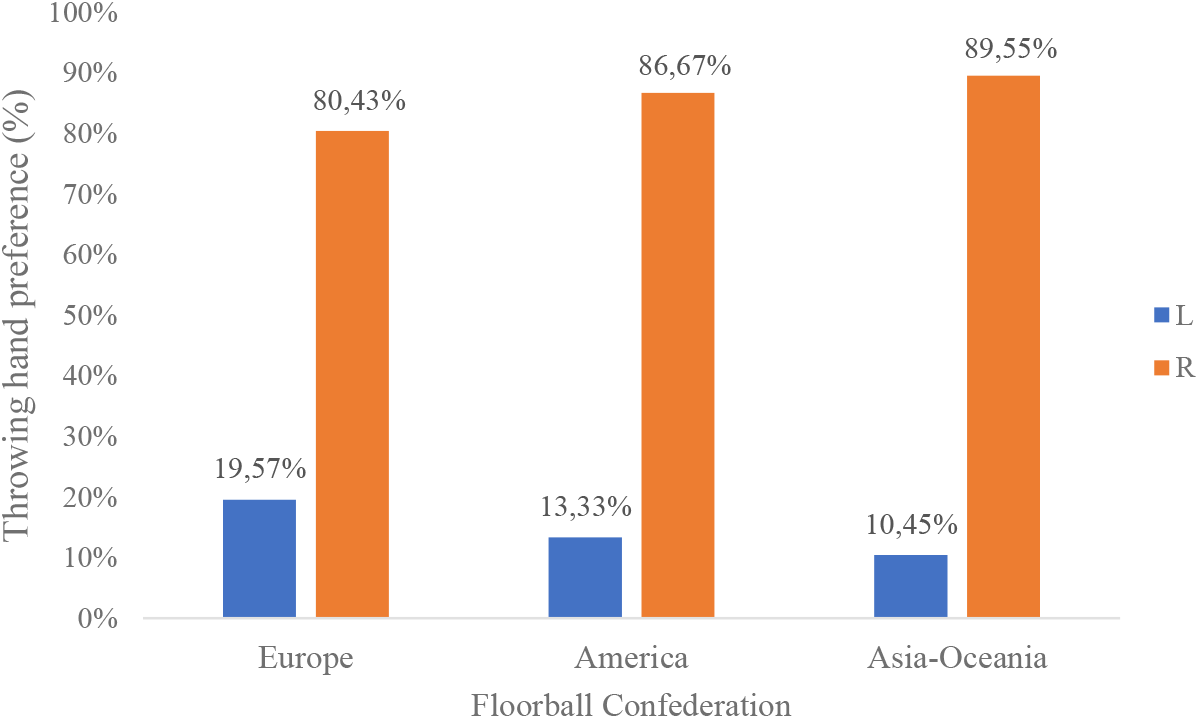
Proportion (%) of left- and right-handed throwing preferences for goalkeepers in each floorball confederation

Sex-dependent variations in goalkeepers’ hand preferences could be identified (Table 1) in that European men were significantly (ꭓ^2^=5.256, p=.022 < .05) more likely to prefer the minority laterality (left-handedness) in comparison to women. The variable and small sample sizes of the Asian-Oceanian and North American cohorts limit further interpretation of the results (n=67 and n=17 respectively).

There were more left-handers among European U19 goalkeepers compared with the corresponding adult sample; however, the difference was not significant (23.7% and 16.7% respectively, ꭓ^2^ = 1.189, *p* = .276 > .05). Also, no significant difference was found among North American or Asian/Oceanian goalkeepers (ꭓ^2^ = .010, *p* = .919 > .05 and ꭓ^2^ = .420, *p* = .517 > .05 respectively), most likely due to the small sample sizes (see Table 1 for details).

## Discussion

As hypothesised, based on the aforementioned anecdotal evidence, there was a complete opposite grip-preference distribution between European and Asian-Oceanian floorball field players. In fact, more than 70% of European players preferred a left-sided grip, whereas almost the same proportion of Asian-Oceanian players preferred the opposite, right-sided grip. For goalkeepers, no such difference was found in the distribution of hand preference between the two confederations. Within the general trend of similarity between the sexes, there was a difference among both the field players and goalkeepers in that male players tended to prefer the confederation-specific minority laterality more often compared to females.

The grip-preference distribution of the European field player sample agrees with previous findings on stick-grip preference in sports where, for example, the majority (67%) of North American ice-hockey players preferred a left-sided shooting grip (13). The study by Puterman et al. suggested that the laterality distribution would have been shaped by a frequency-dependent strategic advantage for right-siders rather than by the players’ general handedness. However, no previously reported evidence could be found to support the geographic variability in lateral motor behaviour between the floorball confederations and, thereby, to explain the opposite grip-side preference in the Asian-Oceanian confederation.

The fact that hand preference among goalkeepers was similar between confederations would indicate that the difference observed among field players was not due to a biological difference or to general handedness. Right-handedness was somewhat more frequent among the Asian-Oceanian goalkeepers (90%) than both their European (80%) and North American (86%) counterparts and, thus, in agreement with the general geographic variabilities in handedness frequency (5).

Men tended to prefer the confederation-specific minority laterality more frequently than women, and this agrees with findings that, in the general population, men are more likely to show a left- or mixed-sided preference (5, 26).

Thus, neither the present findings nor findings from previous studies lend any support to biological factors or general handedness (*individual constraints*) as an explanation for the observed grip-preference distribution difference. One would, in such a case, need to assume that the players in the Asian-Oceanian confederation had developed some kind of opposite specialisation of bimanual motor actions (27, 28). However, the fact that right-handedness is prevalent worldwide (5) in addition to the clear right-handed preference shown in all three goalkeeper samples in the present study would render an opposite handedness highly unlikely. Thus, no genetic variables seem to be able to explain the findings.

Floorball-specific *task constraints* (equipment, game structure and rules applied) are the same across all confederations. Although the sport has developed over a longer time on the European and North American continents compared to the Asian and Oceanian, the current individual game settings for the top level and national team activity are the same. Thus, there should not be any task-specific variations between the sports in each confederation that would have led to an opposite grip preference, which leaves us with *environmental constraints* as the explanation for the inter-confederation grip-preference differences.

One such proposed explanation for the higher frequency of right-sided grip preference in the Asian-Oceanian confederation is the popularity of field hockey in many of the countries (20, 22, 29). Field hockey is a stick and ball game similar to floorball, with the notable difference that players are only allowed to hold the stick with a right-sided grip (30). Therefore, field hockey players transitioning into floorball are assumed to have retained a task-specific (stick and ball manipulating) right-sided laterality. For example, Lai (1999) reported that 35% of the Australian national team in 1998 had no previous floorball experience prior to being recruited (29).

Personal communication with a Swedish former player who is considered one of the first floorball ambassadors on the Asian continent may shed some light on the phenomenon. Together with a few partners, he introduced the sport to Singapore, Malaysia, Indonesia, Myanmar, and Thailand in the early 1990s and later worked to facilitate its development in India, South Korea, Japan, Philippines, New Zealand, and Australia. Interestingly, he did, in fact, play with a right-sided grip but did not believe this to have influenced the grip preference of new players. He explained that several of the first coaches were left-sided, and it was strongly suggested that new players grip the stick with their stronger hand on top (right-hand on a left-sided stick). However, he confirmed that a majority of the initial players and local coaches transitioned to the new sport from field hockey, simultaneously transferring their right-sided grip preferences. Therefore, local coaches introducing the sport to schools encouraged only right-sided playing for younger novices. In addition, according to the above-mentioned floorball ambassador, it so happened that equipment distributors imported only right-sided sticks.

Essentially, the emergence of an opposite floorball-specific grip preference in the Asian-Oceanian field players sample compared with the European and North American samples is believed to be a result of a complex combination of constraints. On a speculative note, could the rather strong presence of field hockey have created an initial right-sided bias for novice floorball players? In this case, the right-sidedness of these first floorball ambassadors could have contributed to the right-sided bias by modelling a right-sided grip pattern. Furthermore, this hypothetical right-sided bias would obviously have benefitted from distributors not importing left-sided sticks and, thereby, might have led to an initial right-sided, sport-specific ecological niche for floorball novices to enter. However, further research is needed to improve the understanding of the sequence in which the different constraining variables have come to create this phenomenon.

The present findings of a complete opposite grip-preference distribution between the European and the Asian-Oceanian confederations would constitute a real-life laboratory for studying how lateral preferences develop within a population. Thus, they afford further studies of the effect an inverted lateral preference might have on several performance variables.

## Conclusion

The present results demonstrate how environmental constraints may influence motor behaviour hand preference, usually believed to be a variable determined in large part by individual (biological) constraints. When the environmental constraints are different, behaviour may take a different path such that the end product is similar but opposite. It would be quite reasonable to believe that this would be the case as well for other behaviours outside sports. We encourage other authors to help explain our findings and to conduct further studies on the topic. The findings would be important for understanding hand preference and, consequently, variables that are dependent on hand preference. In addition, they would add to our understanding of the influence of environmental constraints on motor behaviour and, consequently, for variables dependent on motor behaviour. Lastly, they would increase our understanding of how human behaviour is shaped by the specific conditions under which individuals live.

## Contributors

KEE and AVP conceived the idea and designed the study. KEE collected and analysed the data and interpreted them together with AVP. KEE wrote the first draft of the manuscript and revised it together with AVP. Both authors agreed with the results and conclusions of the manuscript and approved the final version.

## Ethics approval

Not applicable as the study included publicly available data extracted from the internet.

## Data availability statement

The present data were extracted from, and can be found on, the website of the International Floorball Federation (https://floorball.sport/).

